# Deletion of SMC renders FtsK essential in *Corynebacterium glutamicum*

**DOI:** 10.1101/2023.10.14.562338

**Authors:** Feng Peng, Giacomo Giacomelli, Fabian Meyer, Marten Linder, Markus Haak, Christian Rückert-Reed, Manuela Weiß, Jörn Kalinowski, Marc Bramkamp

## Abstract

Structural maintenance of chromosomes (SMC) are ubiquitously distributed proteins involved in chromosome organization. Deletion of *smc* causes severe growth phenotypes in many organisms. Surprisingly, *smc* can be deleted in *Corynebacterium glutamicum*, a member of the Actinomycetota phylum, without any apparent growth phenotype. Earlier work has shown that SMC in *C. glutamicum* is loaded in a ParB-dependent fashion to the chromosome and functions in replichore cohesion. The unexpected absence of a growth phenotype in the *smc* mutant prompted us to screen for unknown synthetic interactions within *C. glutamicum*. Therefore, we generated a high-density Tn-5 library based on wild-type and *smc*-deleted *C. glutamicum* strains. The transposon sequencing (Tn-seq) data revealed that the DNA-translocase FtsK is essential in a *smc* deletion strain. FtsK localized to the septa and cell poles in wild type cells, however deletion of *smc* resulted in a decreased polar FtsK localization. Single-particle tracking analysis further suggests that prolonged FtsK complex activity is both required and sufficient to make up for the absence of SMC, thus achieving efficient chromosome segregation in *C. glutamicum*. Further, single molecule dynamics of FtsK is influenced, albeit indirectly, by DNA-loaded SMC. Deletion of ParB results in an increased of both SMC and FtsK mobility. While the first change agrees with previous data that show how ParB is essential for SMC loading on DNA, the latter suggests that FtsK mobility is affected in cells with defects in chromosome organization. Based on our data we propose a simple, yet efficient mechanism for efficient DNA segregation in *C. glutamicum*, even in absence of SMC proteins.

**Importance:** Faithful DNA segregation is of fundamental importance for life. Bacteria have efficient systems to coordinate chromosome compaction, DNA segregation and cell division. A key factor in DNA compaction is the SMC-complex that is found to be essential in many bacteria. In members of the Actinomycetota *smc* is dispensable, but the reason for this was unclear. We show here that the divisome associated DNA-pump FtsK can compensate SMC loss and the subsequent loss in correct chromosome organization. In cells with distorted chromosomes, FtsK functions for an extended period of time at the septum, until chromosomes are segregated.

## Introduction

The replication and segregation of chromosomes is an essential process in all bacteria and the machineries that are involved in these processes are highly conserved. Bacterial chromosomes usually contain a single origin of replication (*oriC*) and termination site (*ter*). The replication of DNA starts from the *oriC* region and end at the *ter* region (1). The total length of the chromosome exceeds the size of the bacterial cell severalfold and hence, sophisticated packing mechanisms have evolved that help to structure the chromosome to fit the cell compartment. Importantly, despite the dense packing the DNA is still accessible for replication and gene transcription and regulation. Newly replicated chromosomes must be segregated faithfully to the daughter cells in coordination with cytokinesis. Many bacteria employ conserved proteins for these processes. Among the most conserved ones are the structural maintenance of chromosomes (SMC) proteins (2). In many bacteria, SMC is essential for chromosome compaction and segregation (2–4).

SMC proteins are long, rod shaped coiled-coil proteins with a dimerization domain and a ABC-binding cassette head domain. The head domain contains the canonical Walker A and Walker B-motif for ATP binding and hydrolysis (5, 6). The two SMC subunits are associated with a kleisin protein (in bacteria often termed ScpA) that form together a ring-like structure (7). This core structure was shown to contain DNA-binding sites. SMC complexes are DNA-loop extrusion motors that allow DNA-loop formation upon ATP hydrolysis (8). The Kite protein ScpB completes the functional SMC complex and SMC–ScpAB is the prevalent version of SMC protein complexes (9, 10). In *B. subtilis* it was shown that the SMC complex is required for *oriC* segregation and absence of SMC is lethal at fast growth conditions (11, 12). Similarly, *smc* mutations cause cell cycle phenotypes in *Caulobacter crescentus* (*3, 13*). Intestinally, in some bacteria such as *C. glutamicum*, *Mycobacterium smegmatis* and *Staphylococcus aureus* has been reported that SMC can be deleted without induction of a severe phenotype (14–16).

In many bacteria, including firmicutes and actinobacteria, SMC complexes are loaded onto the DNA by a DNA-binding protein ParB (14, 17–20). ParB binds a specific DNA sequence, termed *parS* (21). These ParB binding sites are often clustered around the origin of replication (22). ParB is part of a tripartite DNA segregation system (23). The ParABS system was originally identified as an efficient plasmid segregation system, but it is also encoded on many bacterial chromosomes (24). Deletion of chromosomal ParAB loci often results in severe chromosome segregation and viability phenotypes. In *C. glutamicum* deletion strains of *parA* or *parB* are viable, but lead to a high degree of anucleate cells (25). Chromosome segregation defects also affect correct positioning of the cell division machinery and hence, result in cell length phenotypes (26). After loading by ParB, SMC molecules travel the entire length of the chromosome along the DNA. At least in *B. subtilis* SMC is finally release by XerD at the replication terminus (27).

For *B. subtilis* and *Streptomyces coelicolor* it has been shown that a functional interaction between SMC and the DNA-pump FtsK (or the SpoIIIE homolog of FtsK), exists (28, 29). FtsK is a hexameric membrane anchored protein, that binds DNA by recognition of the FtsK-orienting polar sequences (KOPS) DNA motifs before the end of replication of DNA and transfers chromosomes into two daughter cells (30–32). Upon FtsK/KOPS complex reaching the *dif* site in close proximity to the *ter* region, FtsK recruits and activates XerC-D to resolve chromosome dimers by recombination during replication and finally segregate the chromosome (33). FtsK is part of the mature division machinery and helps to pump remaining DNA through the closing septum to avoid cell division induced DNA damage (34). The nature of the described genetic interaction between FtsK and SMC, however, remained mechanistically unclear.

Using a high-density transposon library in combination with transposon sequencing we revealed that FtsK becomes essential in absence of SMC in *C. glutamicum*. Fluorescence microscopy and single-particle tracing revealed that in absence of SMC, the FtsK DNA-pump remains for an extended time period at the septum, thereby compensating the segregation defect of a *smc* mutant.

## Materials and methods

### Media, bacterial strains, plasmids, and growth conditions

All *C. glutamicum* strains used in this study are based on the parental strain *C. glutamicum* MB001 (35). The strains and plasmids used in this study are listed in Table S1. *E. coli* was grown in lysogeny broth (LB; 10 g/L tryptone, 5 g/L yeast extract and 10 g/L NaCl). *C. glutamicum* was grown in Brain-Heart Infusion Medium (BHI) medium. When needed, kanamycin was added to a final concentration of 30 μg/mL for *E. coli* and 10 μg/mL for *C. glutamicum*.

### Gene deletion, insertion and depletion in *C. glutamicum*

A pK18mobsacB vector was used to delete and insert genes by homologous recombination (36). For gene deletion, regions corresponding to approximately 800 bp upstream and downstream of the target gene were amplified from genomic DNA and cloned into pK18mobsacB. The resulting plasmid was transformed into competent *C. glutamicum* cells. Transformants were spread on BHI agar plate with 10 μg/mL kanamycin at 30°C. The resulting colonies were grown in BHI liquid medium at 30°C for overnight. The culture was then spread on BHI agar plate with 10% sucrose and incubated at 30°C. The deletion strain was picked and checked by colony PCR and for kanamycin-sensitivity by streaking the colonies on BHI and BHI-Kan plates. To insert fluorescence gene sequences into the genome, the respective fluorescence gene sequence was assembled into pK18mobsacB sandwiched between the 800 bp upstream and downstream of the targeted genome locus. Transformation and selection methods were similar to the ones used for gene deletion.

A pSG-dCas9 plasmid was used to construct depletion strains based on the CRISPR/dCas9 system. Briefly, sgRNA was designed via the CHOPCHOP web tool (http://chopchop.cbu.uib.no/). Primers encoding for the forward and reverse ssDNA strands for the sgRNA (Table S2) were annealed by heating them up to 95°C for 4 minutes and gradually decreasing the temperature at a rate of 0.5°C/min until a temperature of 25°C was reached. The resulting sgRNA was ligated using T4 ligase into BsaI digested pSG-dCas9. The resulting plasmid was transformed into *C. glutamicum* competent cells. A concentration of 1 mM IPTG was used to induce the expression of sgRNA and dCas9 to deplete the target gene.

### Tn-5 transposon insertion library construction

Tn-5 transposon insertion library was build based on EZ-Tn5™ <KAN-2>Tnp transposome system (Lucigen, WI, USA). Competent *C. glutamicum* cells were prepared as described previously (37). The transformation was carried out by electroporation of 1 μL of Tn-5 transposome mix into 100 μL competent cell. Transformants were then spread on BHI agar plate with 10 μg/mL kanamycin and growth at 30°C for 16 h. Each library was generated by pooling colonies from five electroporations and consisting of approximately 60,000 transformants.

### Tn-5 transposon library sequencing

Genomic DNA was extracted from strains using NucleoSpin Microbial DNA Mini kit (Macherey-Nagel, Düren, Germany). The preparation of the transposon mutant library for Oxford Nanopore Sequencing has been done as described by Linder et al. (38) with the following modifications: (a). Fragmentation of genomic DNA was done using the Covaris M220 Ultrasonicator to acquire ∼ 750 bp fragments. (b). The Bottom Adapter was expanded to include a 16 bp randomized sequence consisting of four consecutive blocks with the sequence BDHV. The sequence serves the purpose of a unique molecular identifier (UMI) to facilitate the detection of a potential amplification bias. By avoiding one specific base at each position, homopolymers which can cause errors during base calling are avoided. (c) For the first round of PCR, the biotinylated primer was also modified to contain a 10 bp N stretch, acting as an UMI to enhance the detection of amplification biases. This was further tested by performing six separate PCR reactions for MB001 and Δ*smc* each. This was done to monitor a PCR bias arising from sedimented streptavidin beads and to test for “completeness” of the sequencing libraries. Identification of the mapping sites was done using crossalign (https://github.com/MarkusHaak/crossalign). For the purpose of counting the number of transposon insertion sites and mappings, the gene regions were extended 50 nucleotides upstream and shortened by 10 % at the 3’-end to account for insertions in the promoter region(s) and the assumption that disruptions at the 3’-end of the polypetide chain might not be critical for protein function. The resulting counts were normalized using DESeq2 (39) and the log2 fold change for each gene was calculated. In addition, using the normalized counts, the average number of sites per kilobase were calculated for each gene to identify the threshold for classifying a gene as essential as done by Lim et. al. (40)

### Fluorescence microscopy and analysis

Images were obtained using an Axio-Imager M1 fluorescence microscope with an ECPlan Neofluar 100×/1.3 oil Ph3 objective (Carl Zeiss, Jena, Germany). Fluorescence of mCherry and FM-64 was detected using the red channel (EX BP 500/25, BS FT 515, EM BP 535/30). Fluorescence of DNA stained via Hoechst 33342 was detected using the blue channel (EX G 365, BS FT 395, EM BP 445/50). The final concentration of FM-64 and Hoechst 33342 was 10 μg/L.

The regions of interest (ROIs) used to determine fluorescence profiles were drawn manually via the segmented line Fiji tool (line width: 5 pixels – 322.5 nm) and added to the ROI manager. Intensity values for the three channels (Phase, FM4-64 and Hoechst 33342) were extracted with a custom Fiji (41) macro (ProfilingCells_EPI.ijm). Cell length was then automatically calculated based on membrane fluorescence profiles via a custom R (using R Statistical Software (v4.1.2; R Core Team 2021) script (CPAD1). The number of nucleoids and septa was finally determined using a second custom made R script (CPAD2). All the custom-written scripts are available on Github (https://github.com/GiacomoGiacomelli/Cell-Profiles-and-Demographs). R version 1.4.1106, were used for the analysis. R was run via RStudio (42).

For time-lapse experiments, exponentially growing cells were diluted to an OD_600_ of 0.008 in CGXII medium and loaded into a microfluidic chamber (B04A CellASIC, Onix). The temperature of the chamber was maintained at 30°C and the speed of nutrient supply was 0.75 psi. Images were taken in 5 min intervals using an Axio-Observer inverted fluorescence microscope with an ECPlan Neofluar 100×/1.3 oil Ph3 objective (Carl Zeiss, Jena, Germany). The fluorescence of mCherry was detected using the red channel (EX BP 500/25, BS FT 515, EM BP 535/30). To obtain time-resolved data of the cell cycle dependent localisation dynamics of FtsK-mCherry, the custom-made FIJI/R script package Morpoholyzer Generation Tracker was used (43). The extracted fluorescence profiles were further analysed and visualised in Excel.

### Flow cytometry

Flow cytometry was used to analyse the DNA contents per cell. Flow cytometry was performed using a CytoFLEX (Beckman, Coulter, USA) equipped with a 488 nm laser. The method was performed as described before (14). Briefly, cells were fixated in 70% ethanol (1:9 v/v) and washed once in PBS. Cellular DNA was stained using SYBR Green I (Invitrogen, 1:10,000 dilutions) for 15 min. At least 100,000 events were collected per sample at a slow flow rate measuring <5000 events per second. Data analysis was performed using the CytExpert Software 2.4 (Beckman).

### Sample preparation for single-molecule localization microscopy

For single particle tracking, overday cultures of the respective strains (*ftsk-halo,* Δ*smc ftsK-halo,* Δ*parB ftsK-halo,* Δ*smc*Δ*parB ftsK-halo, smc-halo,* Δ*parB smc-halo*) were grown until an approximate OD_600_ of 2 (30°C, 200 rpm). Cells were then stained by addition of HaloTag TMR Ligand (1 ml of cells, 25 nM dye final concentration) and 30 minutes of incubation (30°C, 200 rpm, keep in the dark). Cells were then washed 5 times (4000 rcf, 3 min) with TSEMS and finally resuspended in 500 µl of TSEMS. Freshly prepared cells were immediately loaded on the agarose pad.

For the determination of FtsK-PAmCherry cell levels, overday cultures of the respective strains (*ftsk-pamCherry* and Δ*smc ftsK-pamCherry*) were grown until an approximate OD_600_ of 2.0 (30°C, 200 rpm). Cells were then fixed by adding formaldehyde to a final concentration of 2% followed by incubation for 30 minutes at 30°C and 200 rpm. Cells should be shielded from light both during growth and during the fixation procedure to prevent fluorophores bleaching. After incubation cells were washed three times (3400 rcf, 3 minutes) in 1mL of 1x PBS with 10mM of glycine, incubating for 5 minutes at 30°C and 200 rpm between each wash. Finally, cells were resuspended in 200µL of 1x PBS and 10mM glycine.

### Slides and chambers preparation for single-molecule localization microscopy

Slides and coverslips used for single particle tracking were first cleaned by overnight storage in 1 M KOH, carefully rinsed with ddH_2_O, and subsequently dried with pressurized air. Next, 1% (w/v) low melting agarose (agarose, low gelling temperature, Sigma-Aldrich, Taufkirchen, Germany) was dissolved in medium/buffer TSEMS for *C. glutamicum* for 1 h at 95 °C shaking. Buffer was sterile filtered (0.2 µm pore size) shortly before being used to remove particles. To produce flat, uniform, and reproducible agarose pads, gene frames (Thermo Fisher, Dreieich, Germany) were utilized, and pads were finally allowed to solidify for 1 h at room temperature to be used within the next 3 h.

The µ-Slide 8 Well Glass Bottom chambers used for fixed cells imaging were filled poly-L-lysine (200 µl for each well) and incubated for at least one hour prior the addition of cells to favour cell adherence. Following incubation, poly-L-lysine was removed and the wells were washed three times with 200µL of 1x PBS and 10 mM glycine. The PBS-glycine solution was then removed and 0.5 µl of resuspended (30 s vortex) TetraSpeck Fluorescent Microsphere Standards (0.1 µm diameter – 1:500 dilution of the original stock) were added to each well. 2µL of fixed cells and 200µL 1x PBS and 10mM glycine were also added. Finally, the chambers were centrifuged at 3400 rcf for 3 minutes to allow for the sedimentation of cells.

### Single-molecule localization microscopy

Single-molecule localization microscopy was performed via an Elyra 7 microscope (Zeiss) equipped with two pco.edge sCMOS 4.2 CL HS cameras (PCO AG), connected through a DuoLink (Zeiss), only one of which was used in this study. Cells were observed through an alpha Plan-Apochromat 63x/1.46 Oil Korr M27 Var2 objective in combination with an Optovar 1x (Zeiss) magnification changer, yielding a pixel size of 0.0968 µm. During image acquisition, the focus was maintained with the help of a Definite Focus.2 system (Zeiss). PAmCherry was activated with a 405 nm diode laser (50mW), while fluorescence (both for PAmCherry and Halo-TMR) was excited with a 561 nm diode laser (100 mW). Signals were observed through a multiple beam splitter (405/488/561/641 nm) and laser block filters (405/488/561/641 nm) followed by a Duolink SR QUAD (Zeiss) filter module (secondary beam splitter: LP 560, emission filters: EF BP420-480 + BP495-550).

For single particle tracking, cells were illuminated with the 561 nm laser (60% intensity) in TIRF mode (62° angle). For each time lapse series, 5000 frames were taken with 10 ms exposure time (∼13 ms with transfer time included). The Log Detector of TrackMate v6.0.1 (44), implemented in Fiji 1.53 g (41), was used to identify single particles in the single particle tracking experiments. Sub-pixel localizations were detected for spots characterized by an estimated 0.5 µm diameter and signal to noise ratio threshold of 5. Spots were merged into tracks via the Simple LAP Tracker of TrackMate, with a maximum linking distance of 300 nm and no frame gaps allowed. Only tracks with a minimum length of 5 frames were used for further analysis yielding a minimum of 991 tracks (Table S3).

The phase images of cells were segmented using OUFTI. The combined tracks and cell masks were then imported in the MATLAB package SMTracker (45), which was used for jump distance analysis, MSD determination and the production of averaged cells heat maps. For the determination of FtsK-PAmCherry cell levels, cells were sequentially illuminated with a 405 nm laser and a 561 nm laser.

## Results

### Construction of transposon library and sequencing in *C. glutamicum*

Deletion of *smc* in *C. glutamicum* has no observable growth phenotype under normal growth conditions, despite the fact that replicore cohesion is lost under these conditions (14). Therefore, we wanted to examine how *C. glutamicum* can cope with a grossly altered chromosome organization during fast growth conditions. For this purpose, we constructed high density transposon libraries for wild-type and *smc*-deletion strains using the Tn5-based transposome system (EZ-Tn5™ <KAN-2>), with each library accounting for approximately 60,000 mutants (Figure 1A). Sequencing of the libraries yielded 101,851 unique transposition sites across all samples, of which 40,643 were exclusive for the wild-type strain, and 49,457 were exclusive for the Δ*smc* strain (Figure 1B). Sequencing data revealed that insertions of the transposons in the chromosome are random and cover the entire chromosome (Fig. S1). Further analysis of the insertion frequency confirmed a strong positive correlation between the number of unique insertions within a specific gene and the respective gene length (Figure 1C). This correlation is lost in essential genes, which are characterized by an average insertion rate of 40 per kilobase sequence or lower. For the quantification of essential genes, we excluded genes shorter than 100 bp, along with tRNAs, rRNAs, and insertion elements. We could therefore identify a total of 392 essential genes in the wild-type (13.4 %) and of 377 in the Δ*smc* strain (12.9 %).

**Figure 1:**
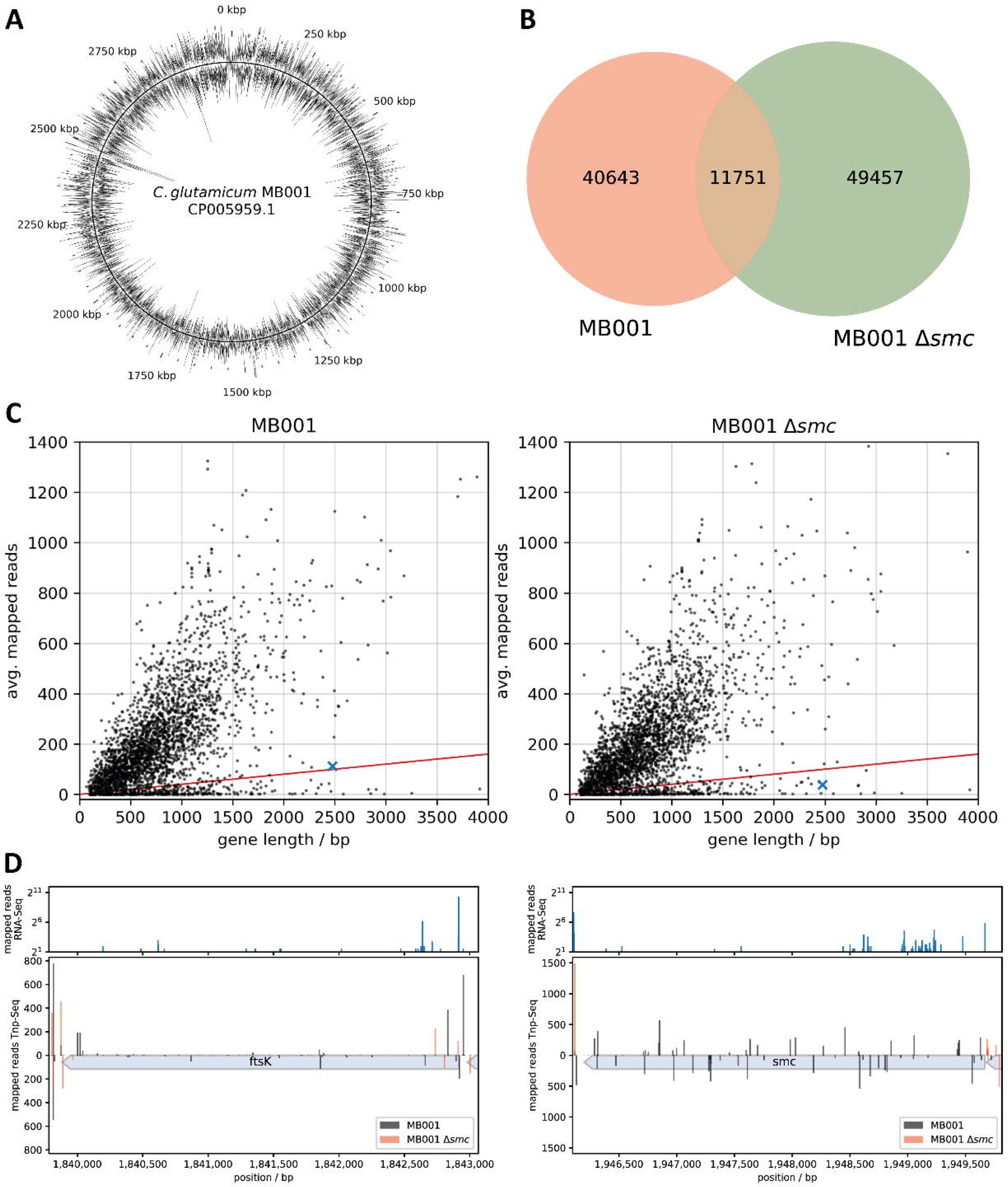
Identification of the genes related to SMC by Tn-seq. (A) Average transposon insertions over 500 bp sequence windows of the wild-type strain mapped to the MB001 genome. (B) Overlap of transposon insertion sites with 10 or more mapped insertions between the two libraries, MB001 wild-type and *Δsmc*. (C) The average normalized number of transposon insertions mapped to every gene and plotted as a function of gene length. Genes with an average rate of 40 transposon insertions per kilobase sequence (red line) are categorized as essential. The *ftsK* gene is highlighted in blue. (D) Raw-counts of transposon insertions mapped to *smc* and *ftsK* in the wild-type (Black) and Δ*smc* (Red) strain. Mapped insertions with reverse orientation of the transposon are displayed as bars below zero. The upper subplots show the counts of the 5’-end enriched RNA-Seq data set from Pfeifer-Sancar et. al. (60) mapped to the MB001 wild-type strain, indicating the position of transcription start sites.

### FtsK is essential in a **Δ***smc* background

We determined synthetic lethality by checking for genes that were characterized by rare or no Tn insertions exclusively in the Δ*smc* genetic background. Most of the genes found to be essential in a *smc* deletion background only are of unknown function (Table 1, Fig. S1), but among those with known activity, we identified *ftsK* (Fig. 1D, Fig. S1). The DNA-pump FtsK functions in DNA segregation during cytokinesis, and deletion of *ftsK* has been reported to lead to DNA damage and segregation defects (32, 46). The number of Tn insertions found in *ftsK* in the Δ*smc* strain is 2.86-fold reduced compared to the wild-type strain. This reduction becomes even more pronounced upon closer inspection, as the majority of insertions in the Δ*smc* strain is located at the 5’-end, upstream of an internal, secondary promoter (Fig. 1D). This is less pronounced in the wildtype, indicating that loss of *ftsK* results in an increased loss of fitness in the Δ*smc* background. The FtsK protein is known to play an essential role in chromosome segregation in *E. coli*. A synthetic interaction of SMC and FtsK has been reported in several organisms including *B. subtilis* and *S. coelicolor*. However, the molecular mechanism of the FtsK SMC interaction remained unclear. It should be noted that we also found ParB among those proteins that become essential in a *smc* deletion strain (Table 1), confirming the close interaction of SMC and ParB for their cellular function.

**Table 1:** Genes essential in Δ*smc*^1^.

| Gene | Description | Log<sub>2</sub> fold change |
| --- | --- | --- |
| cgp_0002 | hypothetical protein | -1,48 |
| cgp_0031 | hypothetical protein | -2,32 |
| cgp_0281 | tRNA-specific adenosine deaminase | -2,84 |
| ccsA | cytochrome c biogenesis membrane protein; DsbD-family | -1,61 |
| cgp_0523 | cytochrome c biogenesis membrane protein; ResB-family | -1,34 |
| menD | 2-oxoglutarate decarboxylase | -2,43 |
| nusG | transcription antitermination protein NusG | -2,03 |
| cgp_0751 | putative membrane protein | -0,96 |
| mrx1 | mycoredoxin 1 | -3,26 |
| fum | fumarate hydratase | -1,75 |
| cgp_1200 | hypothetical protein | -4,01 |
| cgp_1248 | putative GTPase; probably involved in stress response | -0,77 |
| sigE | RNA polymerase sigma factor; ECF-family | -0,65 |
| cgp_1356 | putative rRNA or tRNA methylase | -2,16 |
| cgp_1360 | putative membrane protein | -1,84 |
| cgp_1370 | hypothetical protein | -1,42 |
| cgp_1390 | hypothetical protein | -8,03 |
| cgp_1392 | putative transcriptional regulator; HTH_3-family | -5,83 |
| cgp_1417 | putative acetyltransferase | -3,57 |
| cgp_1433 | hypothetical protein | -2,85 |
| cgp_1491 | hypothetical protein | -6,27 |
| cgp_1618 | hypothetical protein | -2,69 |
| cgp_1669 | putative secreted protein | -1,38 |
| cgp_1766 | putative membrane protein | -0,90 |
| cgp_1792 | putative transcriptional regulator; WhiB-family | -1,13 |
| cgp_1872 | hypothetical protein | -0,84 |
| cgp_2097 | putative DNA or RNA helicase; superfamily II | -0,54 |
| ftsK | DNA segregation ATPase | -1,52 |
| cgp_2159 | hypothetical protein | -2,49 |
| cgp_2340 | ABC-type amino acid transporter; substrate-binding protein | -3,87 |
| cgp_2360 | putative membrane protein | -5,51 |
| fasR | transcriptional regulator of fatty acid synthesis | -2,32 |
| cgp_2850 | hypothetical protein | -1,34 |
| otsB | trehalose phosphatase | -4,46 |
| hpt | hypoxanthine phosphoribosyltransferase | -1,25 |
| cgp_3081 | hypothetical protein | -4,74 |
| cgp_3121 | putative membrane protein | -7,95 |
| parB | putative cell division protein ParB | -0,72 |

<sup>1</sup> Genes not essential in MB001 (> 40 mappings per kb and/or not essential according to Lim et. al. (40)) but essential in $\Delta smc$ and a log<sub>2</sub> fold change <= -0.5

### Depletion of *ftsK* affects cell length and growth in *C. glutamicum*

The low numbers of identified Tn-5 transposon insertions into the *ftsK* gene indicated that FtsK is important in the wild-type strain and essential in the Δ*smc* stain (Fig. 1D). FtsK plays an important role in chromosome segregation during cell division and hence, deletion of *ftsK* has a severe phenotype in most bacteria. In fact, we failed to obtain an *ftsK*-deletion strain in wild type *C. glutamicum*. We therefore used a CRISPR/dCas9 system to deplete the transcription of the *ftsK* gene. To investigate the effect of depletion, we constructed a FtsK-mCherry fusion strain, depleted the protein via the CRISPR/dCas9 system, and analyzed the FtsK-mCherry protein levels by fluorescence microscopy and western blotting. Image analysis and western blotting revealed a significant decrease in FtsK-mCherry signal after depletion (Fig. 2A-B, Fig. S2A). We then combined the *ftsK* depletion strain with the *smc* gene deletion background. Growth analysis of wild-type, Δ*smc* strain and their respective *ftsK*-depletion strains showed that strains with depletion of FtsK grew significantly slower, while wild-type and Δ*smc* strains have similar growth fitness (Fig. 2C). The Δ*smc* strain with simultaneous FtsK depletion displayed the weakest fitness (Fig.2C), indicating the genetic interaction of *smc* and *ftsK* in *C. glutamicum*. Growth analysis in liquid medium supports this notion (Fig. S2C). Similar to the plate-based viability assays, growth in liquid cultures is most affected on the depletion of *ftsK* in a *smc* mutant background. We also observed a severe aggregation phenotype in strains with FtsK depletion in absence of *smc* (Fig. S2C). However, depleting FtsK in the *smc* mutant background resulted in the selection of suppressor mutations, likely in the CRISPR system, so that signal of FtsK-mCherry remained intense at the septum (Fig. S2A).

**Figure 2:**
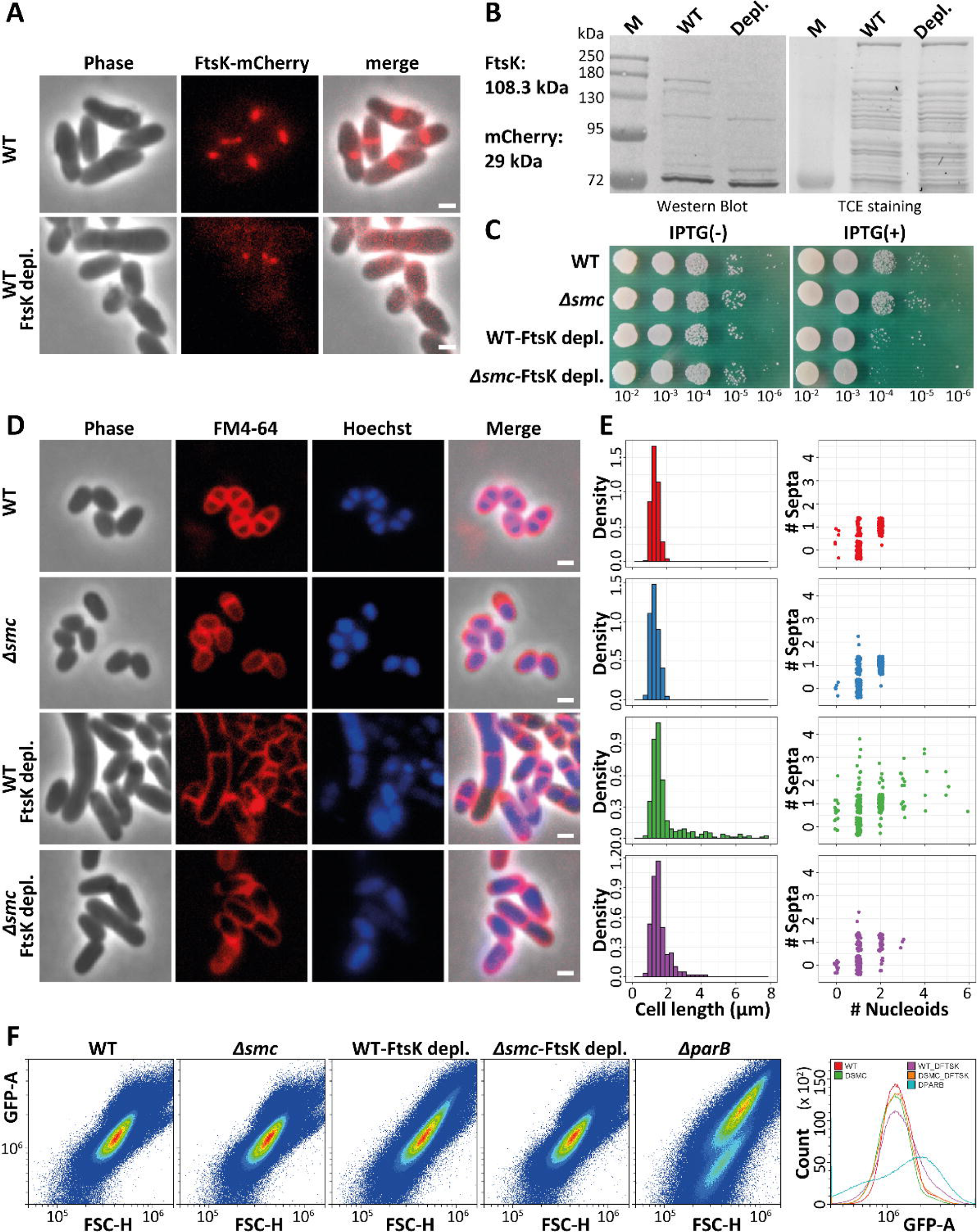
FtsK is essential in an SMC-deleted strain. (A) Microscopic analysis of the strain with depletion of FtsK using CRISPR/ dCas9 system. FtsK fusion with mCherry for detecting the expression of FtsK in wild-type and after FtsK-depletion in wild-type. Protospacer adjacent motif (PAM) site for CRISPR/dCas9 system was at 5’_278-300_3’. Images were obtained at the same exposure time of 1000 ms. Scale bar 2 μm. (B) Analysis of FtsK-mCherry expression using western blotting. TCE stained gels is shown as loading control. (C) Growth fitness analysis of strains with and without inducing the depletion of FtsK. Overnight cultures of wild-type and FtsK-depletion strain were normalized to an OD_600_ of 0.5, serially diluted 10-fold, and spotted (4 μL) onto agar medium with or without 0.1 mM IPTG. (D) Phenotypic anaylsis of strains. Micrographs show cells stained with FM4-64 (Cell membrane, red) and Hoechst (DNA, Blue). The final concentration of FM4-64 and Hoechst was 1 µg/L. Scale bar 2 μm. (E) Analysis of the cell length and the number of septa compare to the number of nucleoids based on microscopic analysis. The number of septa was obtained from the signal peak of FM4-64. The number of nucleoids was microscopically determined after Hoechst staining. (F) Flow cytometry analysis of DNA contents. DNA was stained with SYBR Green I at a final concentration of 1 mM. Analysis was carried out using GFP channel with 200 nm laser. GFP-A represents the signal of SYBR Green I and FSC-H indicates the forward scatter, indicative of cell sizes.

Cell length measurements were performed on cells stained with the membrane dye FM4-64 and nucleoids were stained using Hoechst. As all cell length distributions did not follow normal distributions, (p-value threshold for critical difference: 0.05) (Table S4) we compared the four strains for length via the Kruskal-Wallis multiple comparison test. Cell length distributions comparison revealed that after depletion of FtsK the cell length differed significantly from cells containing native amounts of FtsK for both WT genetic backgrounds while no difference was found between WT and Δ*smc* (Fig. 2D-E). We also counted the number of septa and correlated it with the number of nucleoids as a proxy for determining chromosome segregation/cell division defects (Fig. 2E). The data suggest that *ftsK*-depletion affects the number of septa and nucleoids, and DNA-free cells were also observed (Fig. 2E) (Table S4).

We than used flow cytometry to analyze cell length distribution and DNA content for a larger cell number. These results supported the microscopy data and revealed a clear role for FtsK in correct chromosome segregation (Fig. 2F). Somewhat surprisingly, we observed that depletion of FtsK in the Δ*smc* mutant background resulted in a diminished phenotype compared to the FtsK depletion in wild-type (Fig. 2E). Very likely the synthetic lethality of *smc* and *ftsK* deletions places a strong selection pressure, thereby selecting for suppressor mutations that either reduced the efficient CRISPR interference or generated second site suppressors similar to situations described before (47). Therefore, we turned towards alternative experiments to unravel the functional connection of SMC and FtsK.

### Deletion of SMC influences FtsK localization

To gain insight into the functional interaction of SMC and FtsK, we further analyzed the localization of FtsK-mCherrry in wild-type and Δ*smc* mutant strains. In wild type cells Ftsk was localized at midcell or at the pole (Fig. 3A). In particular, the septal localization was expected due to the role of FtsK in DNA pumping during the late stages of cell division. The clear polar localization was somewhat surprising and may indicate that the component of the *C. glutamicum* divisome remain at the poles before being recruited to the new septum. A similar localization pattern was described for *C*. *crescentus*. In *C. crescentus* FtsK remained at the newly generated cell poles for some time before assembling again at the next septum (48).

**Figure 3:**
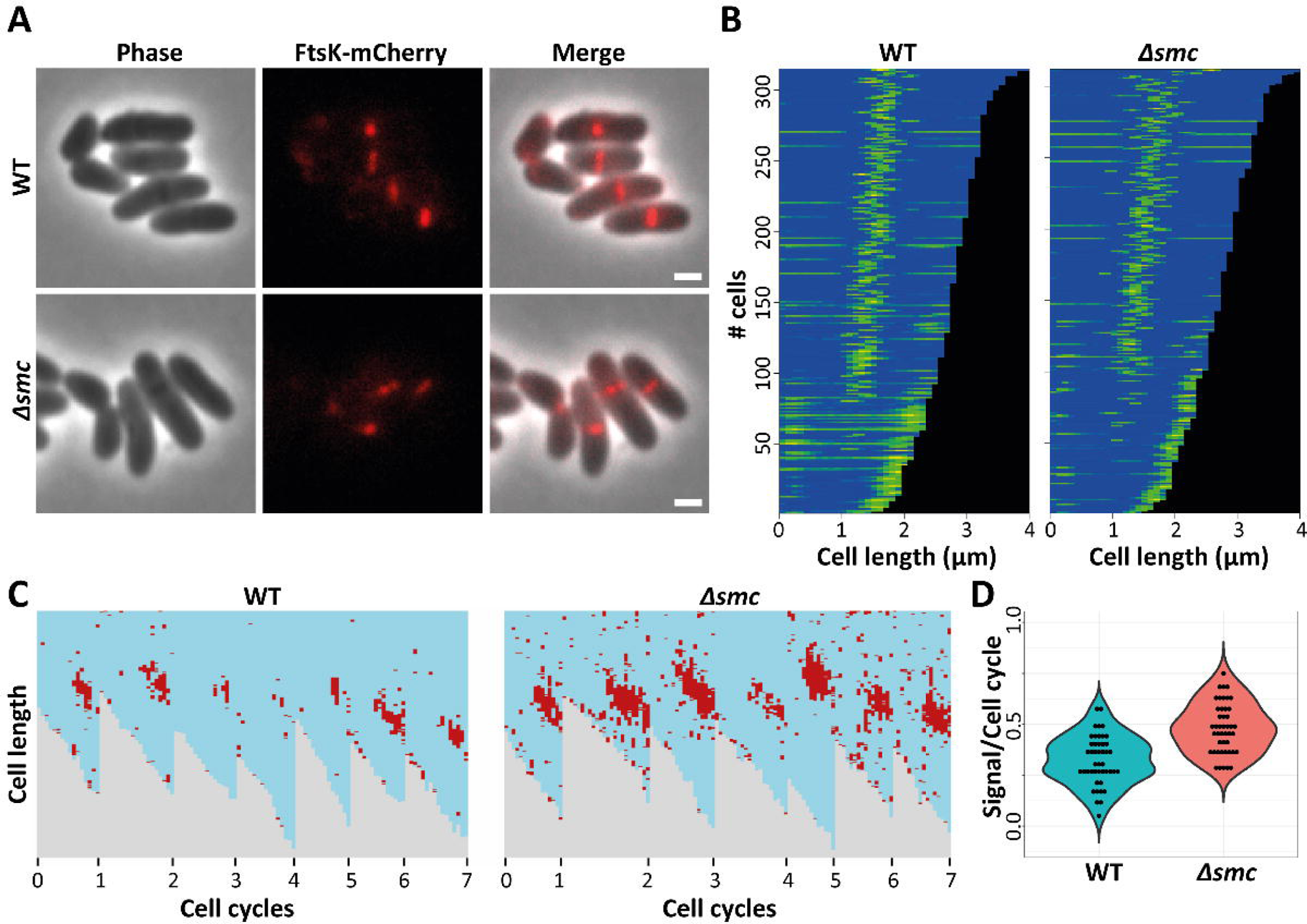
Localization and dynamics of FtsK-mCherry. (A) Cellular localization of FtsK fusion with mCherry. Scale bar 2 μm. (B) Demograph showing the distribution of fluorescence according to cell length. Single-cell fluorescence quantification was performed using Fiji. At least 300 cells were used to generate each demograph. (C) The left panel shows a comparison of the FtsZ-mCherry intensity between the wildtype (WT) and the Δ*smc* genetic background for seven representative cell cycles each, generated form time-lapse micrographs. (D) Statistic evaluation of all 45 measured cell cycles of each genotype reveals a significant difference (Wilcoxon-test; p = 2.1e^-7^). Here the single data points represent the ratio of the presence of the FtsK-mCherry signal versus the duration of the whole cell cycle of each cell.

Interestingly, in pre-divisional cells (cells without a new septum) FtsK localized at both poles in wild-type cells, while the localization was mostly at one pole in the Δ*smc* strain (Fig. 3B). We assume that FtsK remains predominantly at the young cell pole and also in wild type cells we see a more pronounced localization at the younger cell poles. The fact that in Δ*smc* mutant strains the localization of FtsK was almost exclusively at the young cell pole might be a consequence of a slower remodelling of the divisome in these cells. Therefore, we studied the dynamics of FtsK by time-lapse microscopy. In wild type cells FtsK starts localizing at midcell during the last quarter of the cell cycle as judged by monitoring cell growth before division (Fig. 3C). FtsK remains usually until the snapping division at the septum and then largely disperses with some FtsK remaining at the cell poles. Quantification shows that FtsK rarely localized at midcell for more than half of the cell cycle from birth to division. In stark contrast FtsK localization at midcell was significantly enhanced at midcell in the strain lacking *smc*. Here, not only FtsK-mCerry foci appeared brighter at midcell, but also foci remained for a much larger part of the cell cycle (Fig. 3C-D). These data indicate that loss of *smc* leads to a longer localization and functioning of FtsK at the septum. A simple explanation for the enhanced and prolonged FtsK signal at the septum in cells lacking *smc* could be an overexpression of SMC under these conditions. To test this, we used quantitative single molecule localization microscopy (SMLM). A FtsK-PAmCherry was expressed from the native *ftsK* locus in wild type and Δ*smc* strains. SMLM data confirm the localization of FtsK to the septum and to lesser extent also to the cell poles (Fig. S3). Quantification of the events per cell revealed that in wild type and Δ*smc* mutant strains FtsK is present with up to 200 events per cell. The median is around 50 events in wild type and around 70 events Δ*smc* mutant, respectively (Fig. S3). The slight difference is not significant and hence, we conclude that FtsK is not significantly overexpressed when *smc* is deleted.

### Single particle tracking analysis reveals differences in FtsK dynamics in absence of SMC

Single-particle tracking (SPT) is a powerful method to study the dynamic processes in living bacterial cells at nanometer resolution. To visualize the dynamics of FtsK-molecules, we expressed FtsK as a fusion protein with a Halo tag and used TMR dye to track the molecules with SPT (Fig. 4A-B, Fig. S4A). Using the SMTracker program (45), we analyzed the obtained tracks within cells and projected the tracks into a standardized cell with 3 ×1 μm size (Fig. 4B, Fig. S4A). Mapping the tracks revealed that the majority of the FtsK molecules are localized around midcell (Fig. S4A). A heatmap generated from all obtained tracks revealed that the center of the closing septum has the highest density of FtsK molecules (Fig. 4B). In cells lacking *smc* still the majority of confined FtsK tracks localized to midcell (Fig. 4B), however, the heatmap representation revealed that the localization precision to the septum centre was less precise. In general, FtsK occupies a larger and less defined area at midcell in cells deleted for *smc*. From the confinement maps and heatmap of all tracks, it became obvious that the movement of FtsK was less precise around midcell. In addition, to estimate the diffusion constants and relative fractions of FtsK molecules, we performed a jump distance analysis from Squared displacement (SQD) data. The diffusion of FtsK-Halo could be best fitted using three populations (Fig. S4B). About 32.5% of the molecules were present in a confined state in wild-type, while 39.9% were confined in the Δ*smc* strain (Fig. 4A Fig. S4B). The results showed that the confined fraction of FtsK at midcell significantly increased in the Δ*smc* strain.

**Figure 4:**
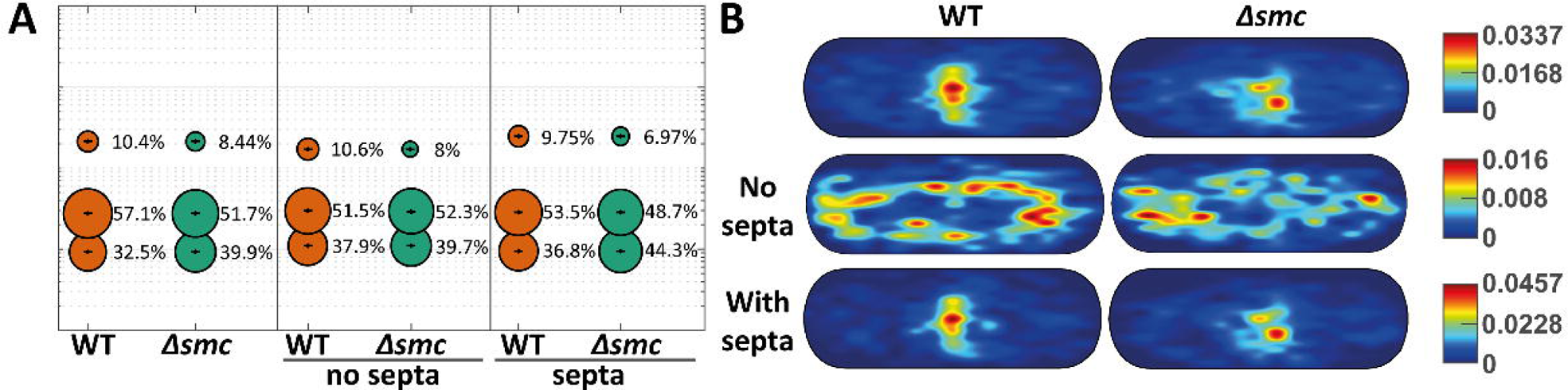
Single-molecule tracking analyses of FtsK in wild-type and *smc*-deleted strains. (A) Summary of the data obtained from SQD analyses. Bubble plot showing single-molecule diffusion rates of the FtsK-Halo fusion protein. Populations were determined by fitting the probability distributions of the frame-to-frame displacement (jump distance) data of all respective tracks to a three components model (fast mobile, slow mobile, and confined protein populations). Separate plots for cell with and without ongoing septation are shown separately (B) Heat map projection of all tracks into a standardized cell of 3 ×1 μm size. Heatmap are shown for all cells, and cells with or without septa.

Since we observed differences in the localization of FtsK at midcell and at the poles in wild-type and Δ*smc* strain, we further analyzed the population of confined molecules of FtsK in cells with and without septa. Interestingly, the confinement maps and heatmap of all tracks showed the difference of FtsK-localization without septa in wild-type and Δ*smc* strain. A small fraction of FtsK molecules were present at both poles in wild-type, while it was present at one pole in Δ*smc* strain (Fig. 4B; Fig. S4A), which was consistent with the of localization analysis of our diffraction limited fluorescence microscopy data. The confinement maps and heatmap also showed the movement of FtsK covered a larger space in the cell with septa in Δ*smc* strain. As expected, the results of jump distance analysed by SQD revealed that the most obvious changes in FtsK dynamics were observed in cells with ongoing septation. Deletion of *smc* led to a reduction of the fast-mobile fraction to about 7% while the confined fraction increased to 44.3% compared to 36.8% in wild type cells (Fig. 4C, Fig. S4B). In addition, our data also revealed that FtsK-molecules have a longer dwell time in Δ*smc* strain background (Table 2, Fig. S5). All these results point towards a longer FtsK function at midcell in the Δ*smc* strain.

**Table 2:**
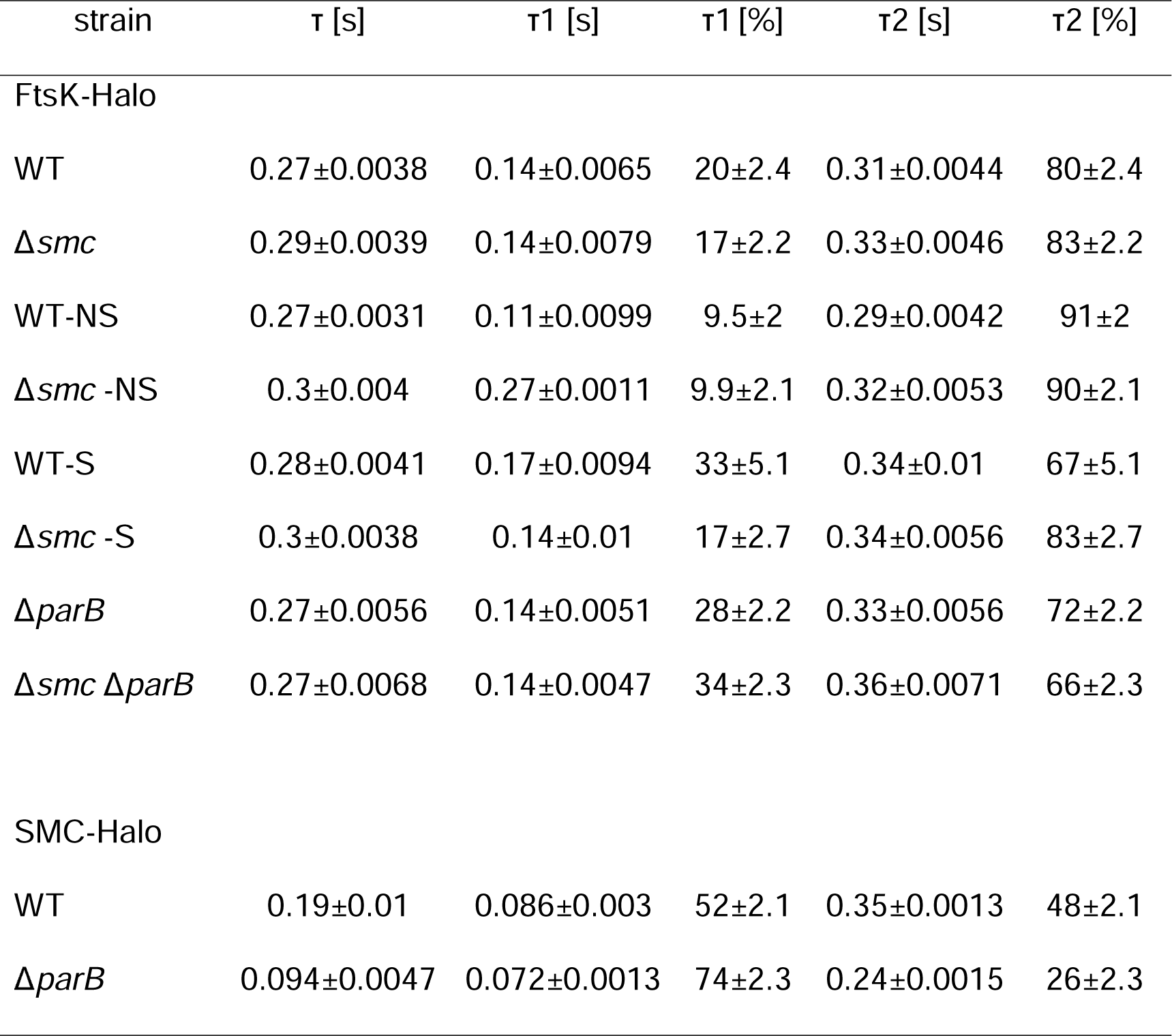
Dwell time of FtsK-molecule and SMC-molecule in different strains.

| strain | $\tau$ [s] | $\tau_1$ [s] | $\tau_1$ [%] | $\tau_2$ [s] | $\tau_2$ [%] |
| --- | --- | --- | --- | --- | --- |
| FtsK-Halo |  |  |  |  |  |
| WT | 0.27±0.0038 | 0.14±0.0065 | 20±2.4 | 0.31±0.0044 | 80±2.4 |
| $\Delta smc$ | 0.29±0.0039 | 0.14±0.0079 | 17±2.2 | 0.33±0.0046 | 83±2.2 |
| WT-NS | 0.27±0.0031 | 0.11±0.0099 | 9.5±2 | 0.29±0.0042 | 91±2 |
| $\Delta smc$-NS | 0.3±0.004 | 0.27±0.0011 | 9.9±2.1 | 0.32±0.0053 | 90±2.1 |
| WT-S | 0.28±0.0041 | 0.17±0.0094 | 33±5.1 | 0.34±0.01 | 67±5.1 |
| $\Delta smc$-S | 0.3±0.0038 | 0.14±0.01 | 17±2.7 | 0.34±0.0056 | 83±2.7 |
| $\Delta parB$ | 0.27±0.0056 | 0.14±0.0051 | 28±2.2 | 0.33±0.0056 | 72±2.2 |
| $\Delta smc \Delta parB$ | 0.27±0.0068 | 0.14±0.0047 | 34±2.3 | 0.36±0.0071 | 66±2.3 |
| SMC-Halo |  |  |  |  |  |
| WT | 0.19±0.01 | 0.086±0.003 | 52±2.1 | 0.35±0.0013 | 48±2.1 |
| $\Delta parB$ | 0.094±0.0047 | 0.072±0.0013 | 74±2.3 | 0.24±0.0015 | 26±2.3 |

### FtsK and SMC dynamics are influenced by ParB

SMC molecules are loaded onto the DNA at the *parS* site by ParB (14, 18, 20). Therefore, the deletion of *parB* leads to a disordered localization of SMC. Deletion of *parB* will therefore affect the movement of SMC and potentially also of FtsK. We analyzed the localization and single-molecule dynamics of SMC using SMC-Halo in wild-type and *parB*-deleted strain backgrounds. SMC localized into smaller clusters in wild-type cells, while we observed a disordered and random localization of SMC in the *parB* mutant cells (Fig. 5A-5B). SPT analysis of SMC revealed that confined and slow diffusive molecules were highly dependent on ParB (Fig. S6A). SMC forms three distinct dynamic populations. About one third (32.1%) of the SMC molecules belong to a confined population in wild-type. About 40% of SMC molecules belong to a slow mobile population, while around 27,9 % of the SMC molecules were highly mobile. The latter likely represents the cytoplasmic fraction of SMC that is not loaded onto DNA. In stark contrast only 5.01% of the SMC molecules were confined in the *parB* deletion strain (Fig. 5D, Fig. S6B). Also, the slow mobile fraction of SMC was decreased to about 17.2% while the vast majority of SMC belong to the high mobile fraction (77.7%). A heatmap presentation also reveals that SMC is far more diffuse localized within the cell when ParB is absent (Fig. 5C). The drastic shift of SMC into the highly mobile fraction is expected when ParB is strictly required for SMC loading in *C. glutamicum*, consistent with earlier findings (14).

**Figure 5:**
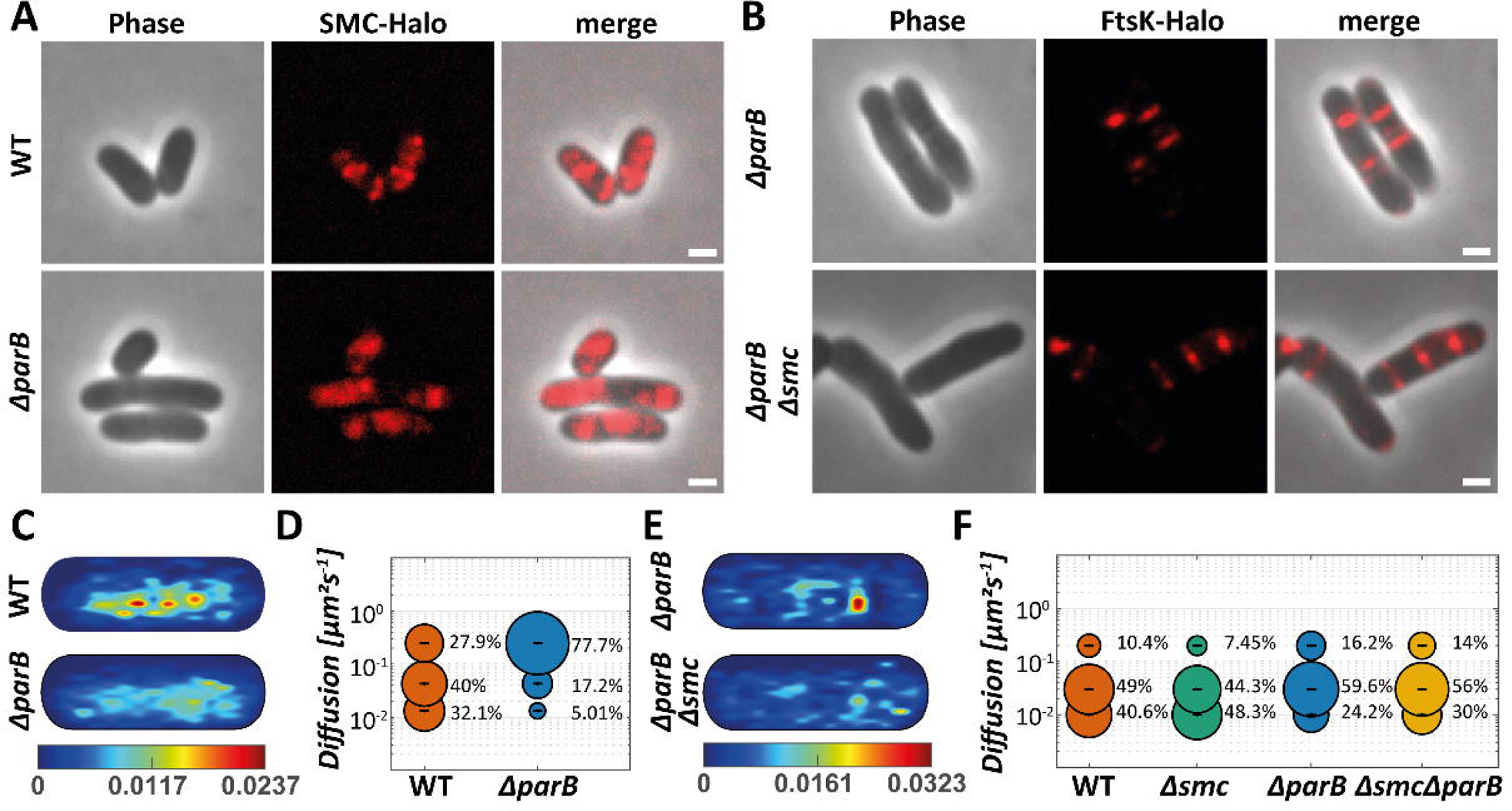
Single-molecule tracking analyses of SMC and FtsK in ParB-deleted strains. (A) Widefield microscopy images of TMR-stained SMC-Halo in the wild-type and ParB-deleted strains. Scale bar 2 μm. (B) Images of FtsK-Halo in ParB-deleted and SMC-ParB deleted strain. Scale bar 2 μm. (C) Heat map projection of all SMC-molecule tracks into a standardized cell of 3 ×1 μm size for wild type and ParB-deleted strains. (D) Bubble plot showing single-molecule diffusion rates of the SMC-Halo fusion protein. Populations were determined by fitting the probability distributions of the frame-to-frame displacement (jump distance) data of all respective tracks to a three components model (fast mobile, slow mobile, and confined protein populations). (E) Heat map projection of all FtsK-molecule tracks into a standardized cell of 3 ×1 μm size in Δ*smc* and Δ*smc*Δ*parB* strains. (F) Bubble plot showing single-molecule diffusion rates of the FtsK-Halo fusion protein in Δ*smc* and Δ*smc*Δ*parB* strains. Populations were determined by fitting the probability distributions of the frame-to-frame displacement (jump distance) data of all respective tracks to a three components model (fast mobile, slow mobile, and confined protein populations).

We next wanted to address whether mobile SMC molecules have an impact on FtsK localization and dynamics. Because we observed major differences in FtsK dynamics only in cells with ongoing septation (Fig. 4), we analyzed of FtsK single particle dynamics in cells with ongoing cell division a *parB* deletion strain and in a double deletion strain, lacking *smc* and *parB* (Fig. 5E-F, Fig. S6C-D). Fluorescence microscopy of FtsK-Halo revealed that the fusion protein still localized as expected mainly to the septa and to the cell poles (Fig. 5B). Consequently, we used the FtsK-Halo fusion construct for SPT analyses. We tracked FtsK-Halo dynamics in wild type, Δ*smc*, Δ*parB*, and the double deletion Δ*smc* Δ*parB*. In wild type cells FtsK dynamics can best be described by three populations. Around 40.6% of the molecules belong to a confined population. We reason that this is the divisome-bound, actively DNA pumping population. A slow mobile fraction of 49% is likely reflecting the membrane-bound, hexameric (or at least oligomeric) structure. A small, fast-diffusive population of around 10% is likely the membrane-bound monomeric fraction of FtsK. In line with the observation described above, deletion of *smc* increases the confined population of FtsK to about 48.3%, indicating that more FtsK is actively engaged in DNA pumping in absence of *smc* (note that we analysed here only cells with ongoing septation) (Fig. 5F, Fig. S6C-D). Interestingly, we observed largely altered FtsK dynamics in absence of *parB*. In Δ*parB* cells the monomeric fraction of FtsK increased to 16.2% while the confined fraction was reduced to 24.2 %. The majority of FtsK in absence of ParB was found in the slow mobile fraction (59.6%). The decrease of the confined fraction is best explained by the reduced septation rate in Δ*parB* cells. Although, we only counted cells with ongoing septation, the ratio between cell length per septum is altered due to cell elongation in Δ*parB* cells. In strains lacking *smc* and *parB* we observed an intermediate phenotype, with around 30% confined FtsK, 56% slow mobile fraction and 14% mobile FtsK (Fig. 5F, Fig. S6D). Heatmap representations reveal that the double deletion of *smc* and *parB* leads to a more dispersed localization of FtsK (Fig. 5E). These data strongly suggest that the active, DNA-pumping fraction of hexameric FtsK increases in absence of SMC, likely compensating for the non-cohesed chromosomes under these conditions.

## Discussion

Using transposon sequencing we identified a synthetic lethal interaction between SMC and FtsK in *C. glutamicum*. The synthetic combination between these two DNA-acting proteins has been described before for *E. coli*, *B. subtilis* and *S. coelicolor* (28, 29, 49). The DNA translocating activity becomes also essential in strains with large chromosomal inversion (50). Thus, DNA organization and FtsK function are closely connected. However, the exact mechanism why FtsK becomes essential in absence of SMC (or other DNA disarrangements) was unclear. FtsK is a DNA pump that helps to translocate the remaining DNA strand trough the closing septum during cytokinesis (31, 32). SMC is required for correct folding and compaction of the DNA (8, 14, 18, 20). In a simplistic model it seems therefore clear that the chromosomes are less well segregated in absence of SMC and hence, more DNA might still be entrapped in the wrong half of the newly forming daughter cells during septation. If this DNA would not be entirely translocated in the correct cell half, cytokinesis would guillotine the DNA and lead to loss of complete chromosomes in daughter cells, as seen in *B. subtilis* SMC mutants (12). This could be prevented by a) longer working of the DNA pump FtsK, b) a more efficient (accelerated) transport process, or c) a delay in cytokinesis. In order to distinguish between these possibilities, we have performed time lapse imaging and single particle tracking analyses. Because of its important role in DNA segregation and cytokinesis FtsK mutations have severe phenotypes. In *E. coli* a loss of FtsK can be compensated by a suppressor mutation in FtsA (51). Although our Tn-Seq screen identified several transposon insertions into the FtsK gene in wild type background, we were unable to construct a clean deletion of the gene. This may in part be influenced by the fact that gene deletions are carried out via a two-step recombination and the second recombination results in the mutant or in a reversion back to wild type. If the mutant has a strong phenotype it is not unusual to select only for revertants. Therefore, it is somewhat challenging to analyze the synthetic effects of *smc* and *ftsK* mutations. To confirm that a simultaneous deletion of *smc* and *ftsK* results in an increased growth phenotype, we constructed a FtsK depletion strain using a CRISRi approach. Viability assays show that the double mutant is indeed less viable, supporting the Tn-Seq results.

The subsequent analysis clearly supports the notion that FtsK remains longer active at the site of division in absence of SMC. Time lapse imaging showed that FtsK is present at midcell for a short duration in wild type cells, while the protein remained much longer at the septum in cells lacking SMC. A model in which FtsK is longer active at the septum to allow complete segregation of the DNA is also supported by SPT data. In absence of SMC we found that FtsK is in general less dynamic. In particular the fast-mobile fraction was found to be reduced. Single particle tracking data for FtsK proteins in vivo are scarce. The only single molecule tracking data available so far stem from the SpoIIIE protein in *B. subtilis* (52). However, SpoIIIE was shown to act mainly at the sporulation septum and is mainly localized evenly in the membrane during vegetative growth of *B. subtilis* (52, 53). For SpoIIIE two main populations with diffusion constants of 0.34 µm^2^ s^-1^ and 0.096 µm^2^ s^-1^ have been reported (52). These values are very similar to the diffusion constants that we have observed for FtsK in *C. glutamicum* (D*_Fast_* = 0.394 ± 0.002 (μm^2^s^-1^), D*_Slow_* = 0.0509 ± 0 (μm^2^s^-1^)). It was concluded that the slower SpoIIIE fraction corresponds to the hexameric form and the fast fraction to the unassembled (either monomeric or trimeric) form of the FtsK-ATPase (52). Importantly, we observed here additionally a more confined fraction of the FtsK molecules (D*_statict_* = 0.017 ± 0.002 (μm^2^s^-1^)). This fraction is mainly found at active septa and likely shows the divisome associated FtsK population. We found that the dynamics of FtsK in *C. glutamicum* is altered in cells lacking SMC or ParB, however in opposite directions. While absence of SMC leads to a decrease of the mobile fraction and an increase in the confined population, deletion of ParB leads to a significant mobilization of FtsK molecules. Deletion of ParB gives rise to a larger percentage of DNA-free minicells. This may lead in many cases to little or no DNA entrapped within a septum. Under these conditions FtsK might therefore be less confined. This finding indicates that apparently, FtsK molecules are only confined when there is DNA to be segregated in a closing septum.

We also determined the molecular dynamics of SMC in *C. glutamicum*. SMC is loaded to DNA on *parS* sites with the help of ParB proteins (14). CTP loaded ParB binds to *parS* sites, thereby closing a ring-like structure to allow sliding of the ParB dimers along DNA to sites flanking *parS*. Unloading of ParB occurs to slow CTP hydrolysis. Corynebacterial ParB undergoes a liquid-liquid phase separation upon CTP and *parS* addition (54). Recruitment of SMC and topological organization of the origin region might be influenced by phase separation. A CTP hydrolysis mutant, ParB175A, is unable to phase separate and in vivo this mutation leads to stable tethering of SMC to the *parS* sites (14, 54). In wild type conditions SMC, once loaded, migrates slowly across the entire nucleoid, until it is released at the terminus region by interaction with XerCD (27, 55). In many bacteria including *C. glutamicum* this SMC dynamics leads to replichore cohesion (14). In vitro analysis with purified SMC complexes from different organisms shows that these proteins are loop-extrusion machines (56, 57). Loop extrusion and compacting the circular bacterial chromosome by SMC are likely important for correct segregation of the chromosomes during replication and division (8). It is therefore astonishing that only a few dozen molecules of SMC (in *B. subtilis* estimations of around 30 SMC dimers were reported) are present in vivo (58). In *B. subtilis* two populations of SMC dynamics have been described. A fast-diffusive fraction of about 52% (diffusion coefficient 0.53 µm^2^ s^-1^) and a slow diffusive fraction of about 48 % (diffusion coefficient 0.1 µm^2^ s^-1^) was identified (59). These values compare well with the data that we obtained for *C. glutamicum* (D*_Fast_* = 0.453 ± 0.001 (μm^2^s^-1^), D*_Slow_* = 0.071 ±

0.001 (μm^2^s^-1^)). Importantly, we see also a confined fraction of about 32 % of the SMC molecules in wild type cells (D*_Static_*= 0.023 ± 0 (μm^2^s^-1^)). Confined SMC molecules tend to cluster and likely these represent areas in the cell where SMC is loaded onto the DNA (e.g. the *oriC* region). Deletion of ParB leads to a drastic decrease of the confined fraction to only around 5%, while the slow and fast mobile fractions increase to 17 % and 77.7 %, respectively. These data indicate that SMC is unable to bind DNA efficiently in absence of ParB in vivo. Since we observed an increase in FtsK mobility in absence of ParB, a fast diffusive, unbound SMC could directly trigger FtsK dynamics. To test this, we used a double *smc parB* mutant. In these conditions, FtsK remains equally mobile comparted to the *parB* single mutant, ruling out that unloaded SMC might have a direct effect on FtsK dynamics.

Our data are in line with the hypothesis that FtsK dynamics in vivo depends mainly on the cargo DNA. Once FtsK is successfully recruited to the site of division and after loading onto DNA, it keeps actively pumping DNA until the entire chromosome is segregated. In case needed, cell division is postponed until this task has been achieved. However, FtsK seems to be more mobile if there is less or no DNA to pump as in case of the ParB mutant. The reason why we see opposing effects on FtsK dynamics in a SMC and ParB mutant is that only the ParB mutation gives rise to a severe mis-segregation phenotype due to the loss of directed segregation of the origin region. The SMC phenotype is loss in replichore cohesion, but this is not translated into a segregation phenotype, which consequently has little effect on cell growth. Only when FtsK is additionally inactivated the *smc* deletion becomes lethal. Finally, our data also reveal that ParB becomes essential in a Δ*smc* strain background. ParB is essential for origin segregation in *C. glutamicum* and lack of a directed segregation of newly replicated origins becomes lethal when the chromosome is simultaneously imperfectly compacted.

