## Supplemental Material for "Deletion of SMC renders FtsK essential in *Corynebacterium glutamicum*"

### Supplementary Data Legends

**Figure S1**: (separate pdf) Map of *Corynebacterium glutamicum* MB001 with transposon insertion sites and average number of mappings per 500 bp sliding window (100 bp step size). Triangles in black (above) indicate TNP sites with >= 10 mappings in MB001, triangles in red (below) mark TNP sites in Δ*smc*, irrespective of the strand. Lines denote the average number of mappings per 500 bp sliding window (100 bp step size), split by strand. Genes marked orange/green/blue are classified as essential (< 40 mappings per kb) in Δ*smc*/MB001/both strains, while those in grey are considered non-essential. Genes in white were not classified.

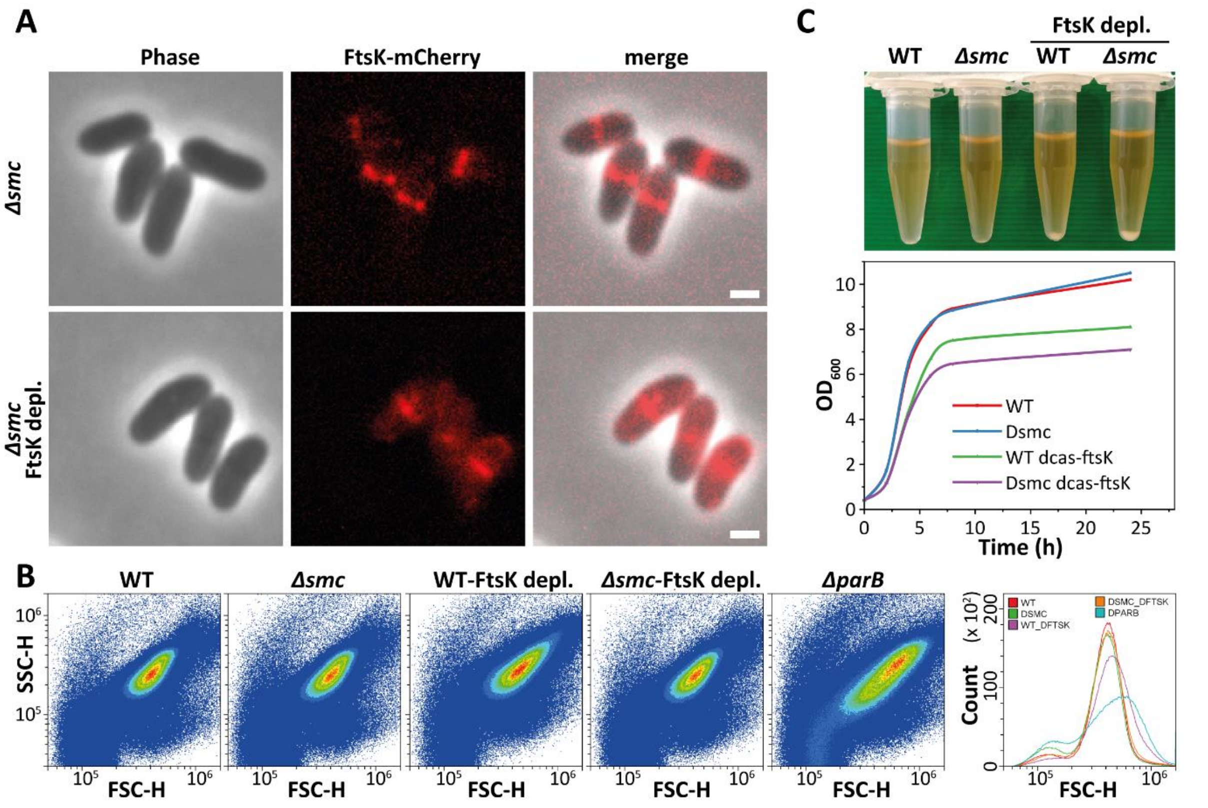

**Figure S2:** (A) Micrographs showing the localization of FtsK-mCherry SMC-deleted strain (upper panel) and with CRISPRi depleted FtsK (lower panel). Scale bar 2 µm. (B) Flow cytometry analysis of cell length using FSC-H as a measure of cell length. (C) Growth analysis of strains with depletion of FtsK. Overnight cultures of wild-type and FtsK-depletion strains were normalized to an $OD_{600}$ of 0.5 in new BHI medium, then $OD_{600}$ were taken every two hours. IPTG was added into the medium at 1h with a final concentration of 0.1 mM. Note that the cells with induced FtsK depletion started to aggregate and sediment in a test tube (upper panel). This effect was increased in cells simultaneously carrying a *smc* deletion.

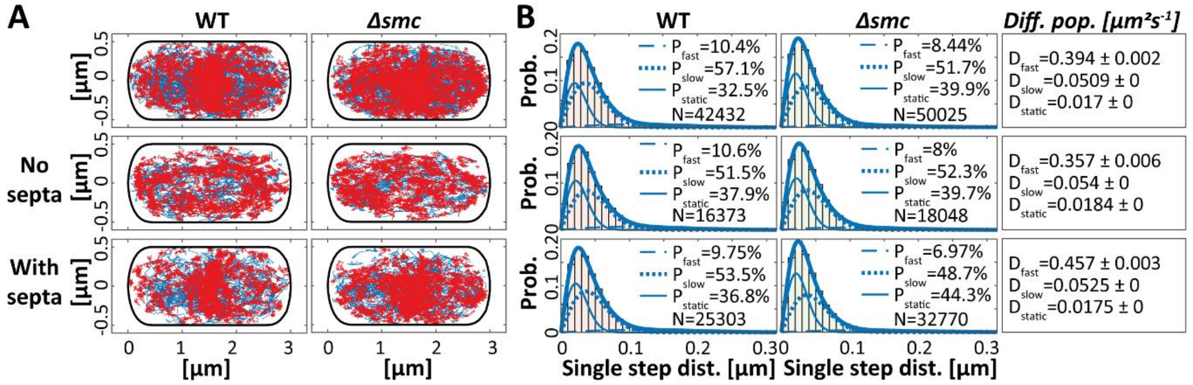

**Figure S3**: (A) Single-molecule tracking analyses of FtsK-Halo in wild-type and *smc*-deleted strains. Projection of all tracks into a standardized cell of 3 ×1 µm size. Tracks moving within a radius of 97 nm for at least 8 steps are shown in red. Tracks moving less confined (faster) are shown in blue. (B) Populations of protein dynamics were determined by fitting the probability distributions of the frame-to-frame displacement (jump distance) data of all respective tracks to a three-component model (fast mobile, slow mobile,and confined protein populations).

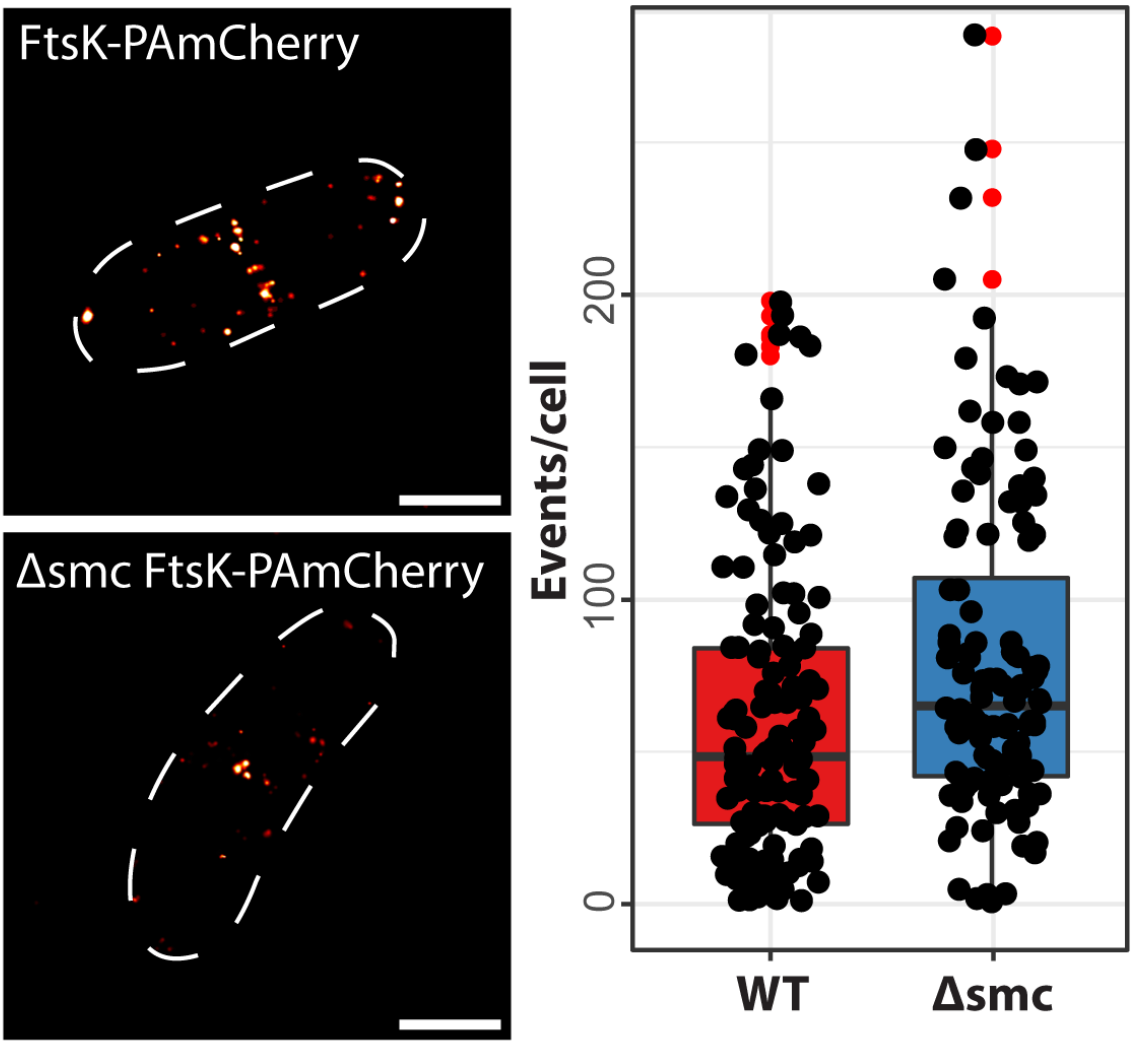

**Figure S4**: PALM analysis of FtsK localization. FtsK-PAmCherry is expressed from its native locus in wild type and Δ*smc* cell backgrounds. Quantification of events for FtsK-PAmCherry in both strain backgrounds.

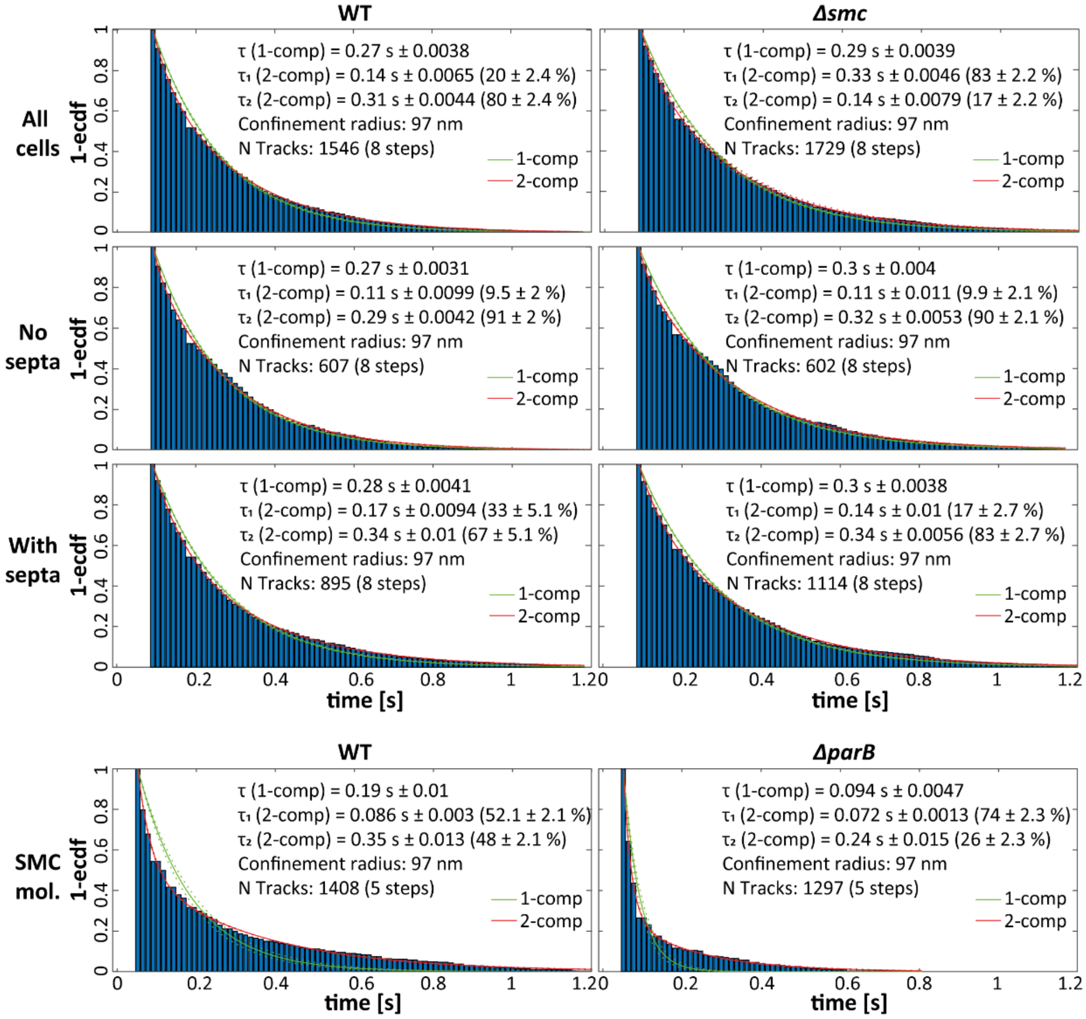

**Figure S5**: Dwell time analysis of FtsK-Halo (upper panels) and SMC-Halo (lower panel). Determination of average dwell times of the indicated fusion proteins via a double exponential decay fit to the survival function (probability of molecules being confined for at least a certain amount of time), here fitted with a one and two component model (green and red lines, respectively). Dwell time of FtsK-Halo molecules in wild-type (left) and SMC-deletion strain (right). Dwell time of SMC-molecules in wild-type (left) and a ParB-deletion strain (right).

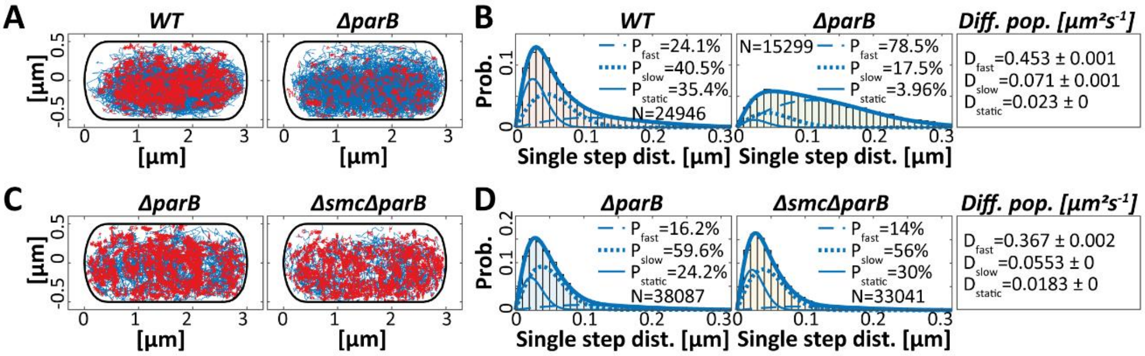

**Figure S6**: Single-molecule tracking analyses of SMC-Halo and FtsK-Halo in different strain backgrounds. (A) Confinement maps of all SMC-molecule tracks in cells of wild-type and ParB-deleted strain. (B) Analysis of the Jump distance using squared displacement analyses (SQD) to estimate the diffusion constants of three diffusive states. (C) Confinement maps of all FtsK-Halo molecule tracks in ∆*parB* and ∆*parB* ∆*smc* strains. (D) Analysis of the Jump distance using squared displacement analyses (SQD) to estimate the diffusion constants of three diffusive states.

**Table S1: Strains and plasmids used in this study.**

| Strains or plasmids | Description | Source or reference |
|---|---|---|
| strains | | |
| *E. coli* DH5α | F- *thi-1 endA1 hsdR17*(r- m+) *supE44* Δ*lacU169* (φ80*lacZ_M15*) *recA1 gyrA96 relA1*; host for cloning procedures | NEB |
| MB001 | ATCC 13032 DCGP1 (cg1507-cg1524), DCGP2 (cg1746-cg1752), DCGP3 (cg1890-cg2071) | (Baumgart et al., 2013) |
| Δ*smc* | MB001 deleted of *smc* | This study |
| Δ*parB* | MB001 deleted of *parB* | This study |
| Δ*smc*Δ*parB* | MB001 deleted of *smc* and *parB* | This study |
| FtsK-mCherry | Fusion FtsK with mCherry in MB001 wild-type | This study |
| Δ*smc* FtsK-mCherry | Fusion FtsK with mCherry in Δ*smc* | This study |
| Ftsk-halo | Fusion FtsK with Halo tag in MB001 wild-type | This study |
| Δ*smc* FtsK-halo | Fusion FtsK with Halo tag in Δ*smc* | This study |
| Δ*parB* FtsK-halo | Fusion FtsK with Halo tag in Δ*parB* | This study |
| Δ*smc*Δ*parB* FtsK-halo | Fusion FtsK with Halo tag in Δ*smcparB* | This study |
| SMC-halo | Fusion SMC with Halo tag in MB001 wild-type | This study |
| Δ*parB* *smc*-halo | Fusion SMC with Halo tag in Δ*parB* | This study |
| FtsK-PAmCherry | Fusion FtsK with PAmCherry in MB001 wild type | This study |

| | | |
|---|---|---|
| Δ*smc* FtsK-PAmCherry | Fusion FtsK with PAmCherry in Δ*smc* | This study |
| FtsK-depletion | Depletion of FtsK in MB001 wild-type | This study |
| Δ*smc* FtsK-depletion | Depletion of FtsK in Δ*smc* | This study |
| FtsK-mCherry FtsK-depletion | Depletion of FtsK in FtsK-mCherry | This study |
| Δ*smc* FtsK-mCherry | Depletion of FtsK in Δ*smc* FtsK-mCherry | This study |
| Plasmids | | |
| pK18*mobsacB* | Kan, vector for allelic exchange in *C. glutamicum* | (Schäfer et al. 1994) |
| pK18*mobsacB*-*smc* | Kan, pK18*mobsacB* with upstream and downstream of *smc* | This study |
| pK18*mobsacB*-*parB* | Kan, pK18*mobsacB* with upstream and downstream of *parB* | This study |
| pK18*mobsacB*-*ftsK-mcherry* | Kan, pK18*mobsacB* for inserting of *mCherry* into downstream of *ftsK* | This study |
| pK18*mobsacB*-*ftsK-halo* | Kan, pK18*mobsacB* for inserting of *halo* into downstream of *ftsK* | This study |
| pK18*mobsacB*-*smc-halo* | Kan, pK18*mobsacB* for inserting of *halo* into downstream of *smc* | This study |

| | | |
|---|---|---|
| pSG-dCas9 | Kan, vector for depletion of target gene in *C. glutamicum* | Bai lab |
| pSG-dCas9-*ftsK* | Kan, vector for depletion of *ftsK* | This study |

**Table S2: Primers used in this study for the construction of sgRNA.**

| Primers | Sequence | Source or reference |
|---|---|---|
| ftsksg3F | TGTGGACGACGACCAAGTTCACCAAG | This study |
| ftsksg3R | AAAACTTGGTGAACTTGGTCGTCGTC | This study |

**Table S3: The number of cells and tracks used in this study**.

| Strain | Cells | Tracks | Average cell length | Tracks/cell |
|---|---|---|---|---|
| FtsK-Halo | | | | |
| WT | 79 | 2795 | 2.79 | 35.7 |
| Δ*smc* | 106 | 2772 | 2.54 | 25.8 |
| WT-NS | 42 | 1063 | 2.26 | 25.0 |
| Δ*smc*-NS | 43 | 991 | 2.12 | 22.7 |
| WT-S | 48 | 1621 | 2.78 | 33.8 |
| Δ*smc*-S | 59 | 1759 | 2.71 | 29.2 |
| Δ*parB* | 41 | 2594 | 5.46 | 59.8 |
| Δ*smc*Δ*parB* | 92 | 2272 | 5.73 | 25.8 |
| SMC-halo | | | | |
| WT | 86 | 1803 | 2.94 | 21.7 |

| | | | | |
|---|---|---|---|---|
| Δ*parB* | 64 | 1695 | 4.25 | 25.8 |

**Table S4: Statistics values**

| Shapiro Test for Normality – shapiro.test() | | | | |
|---|---|---|---|---|
| Strain/s | Cells | Parameter | p.value | Figure |
| MB001 | 284 | Length | 9.26e-05 | 2 |
| Δ*smc* | 333 | Length | 1.16e-06 | 2 |
| FtsK-depletion | 312 | Length | < 2.2e-16 | 2 |
| Δ*smc* FtsK-depletion | 233 | Length | 1.263e-14 | 2 |
| MB001 | 284 | Blue Peaks | | |
| Δ*smc* | 333 | Blue Peaks | | |
| FtsK-depletion | 312 | Blue Peaks | | |
| Δ*smc* FtsK-depletion | 233 | Blue Peaks | | |
| MB001 | 284 | Red Peaks | | |
| Δ*smc* | 333 | Red Peaks | | |
| FtsK-depletion | 312 | Red Peaks | | |
| Δ*smc* FtsK-depletion | 233 | Red Peaks | | |

| Multiple Comparison Kruskal Wallis - krukalmc() - p-value 0.05 | | | | |
|---|---|---|---|---|
| Comparison (Par.) | Obs. Dif. | Critical Dif. | Difference | Figure |
| MB001 / Δ*smc* (Length) | 21.7 | 71.5 | FALSE | 2 |
| MB001 / FtsK-dep. (Length) | 257.2 | 72.6 | TRUE | 2 |

| | | | | |
|---|---|---|---|---|
| MB001 / Δ*smc* FtsK-dep. (Length) | 185.4 | 78.3 | TRUE | 2 |
| Δ*smc* / FtsK-dep. (Length) | 278.9 | 69.8 | TRUE | 2 |
| Δ*smc* / Δ*smc* FtsK-dep. (Length) | 207.1 | 75.6 | TRUE | 2 |
| FtsK-depletion / Δ*smc* FtsK-dep. (Length) | 71.8 | 76.7 | FALSE | 2 |
