## Supplementary figures and images for "Deletion of SMC renders FtsK essential in *Corynebacterium glutamicum*"

### Figure S1

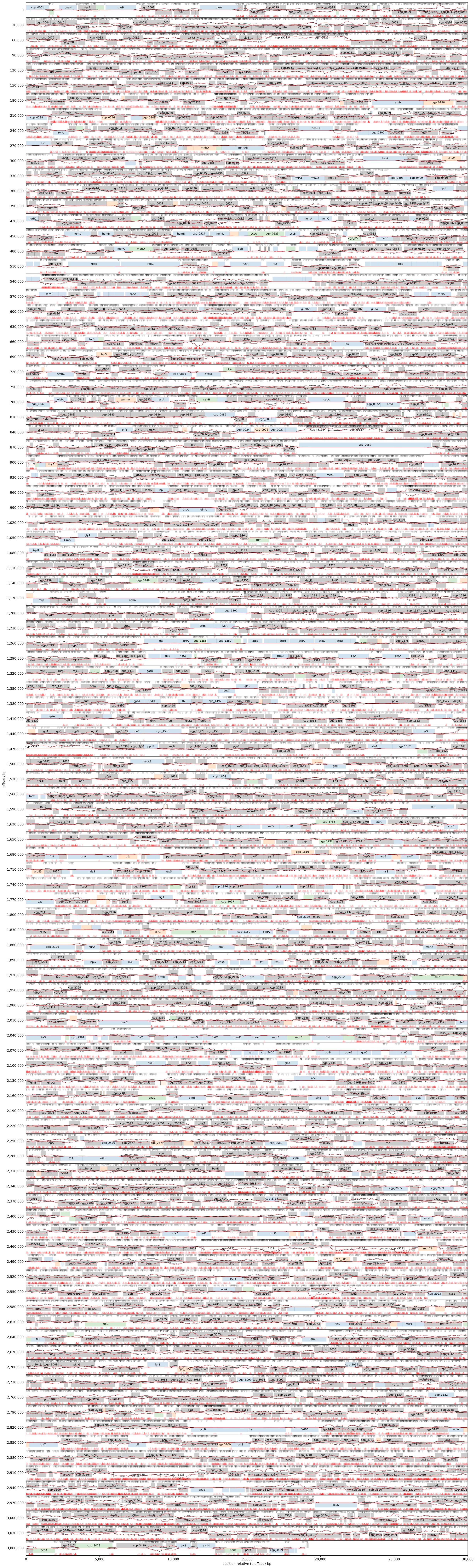
